# Reduced visitation to the buzz-pollinated *Cyanella hyacinthoides* in the presence of other pollen sources in the hyperdiverse Cape Floristic Region

**DOI:** 10.1101/2021.04.17.440253

**Authors:** Jurene E. Kemp, Francismeire J. Telles, Mario Vallejo-Marin

## Abstract

Many plant species have floral morphologies that restrict access to floral resources, such as pollen or nectar, and only a subset of floral visitors can perform the complex handling behaviours required to extract restricted resources. Due to the time and energy required to extract resources from morphologically complex flowers, these plant species potentially compete for pollinators with co-flowering plants that have more easily accessible resources. A widespread floral mechanism restricting access to pollen is the presence of tubular anthers that open through small pores or slits (poricidal anthers). Some bees have evolved the capacity to remove pollen from poricidal anthers using vibrations, giving rise to the phenomenon of buzz-pollination. These bee vibrations that are produced for pollen extraction are presumably energetically costly, and to date, few studies have investigated whether buzz-pollinated flowers may be at a disadvantage when competing for pollinators’ attention with plant species that present unrestricted pollen resources. Here, we studied *Cyanella hyacinthoides* (Tecophilaeaceae), a geophyte with poricidal anthers in the hyperdiverse Cape Floristic Region of South Africa, to assess how the composition and relative abundance of flowers with easily accessible pollen affect bee visitation to a buzz-pollinated plant. We found that the number of pollinator species was not influenced by community composition. However, visitation rates to *C. hyacinthoides* were negatively related to the abundance of flowers with more accessible resources. Visitation rates were strongly associated with petal colour, showing that flower colour is important in mediating these interactions. We conclude that buzz-pollinated plants might be at a competitive disadvantage when many easily accessible pollen sources are available, particularly when competitor species share its floral signals.

## Introduction

The majority of flowering plants are pollinated by animals (Ollerton et al. 2011), and most animal-pollinated species offer resources such as pollen, nectar, oils and scents as rewards to attract floral visitors. The degree to which these resources are accessible to floral visitors varies between plant species, and the accessibility of resources is often modulated through morphological restrictions, such as nectar tubes or keel flowers (Córdoba and Cocucci 2011, Santamaría and Rodríguez-Gironés 2015). Although morphological restrictions can limit or prevent inefficient pollen vectors or resource thieves from gaining access to floral resources (van der Kooi et al. 2021), these barriers can also influence visitation by efficient pollinators, particularly when floral resources can more easily be obtained from flowers that do not restrict access to resources.

Flowers with morphologies that require complex handling behaviours for resource extraction co-occur and potentially compete with flowers that offer more easily accessible resources. Floral visitors that have the ability to extract resources from morphologically complex flowers may preferentially visit complex flowers either when these flowers offer larger resource quantities or higher quality resources than flowers with unrestricted resources (Arroyo and Dafni 1995, Warren and Diaz 2001), or when the probability of obtaining resources is higher in complex flowers because few other species can access the resources (Warren and Diaz 2001). Alternately, if flowers with more easily accessible resources are abundant in a community, the costs associated with learning to handle complex flowers, as well as the time and energy costs of foraging on complex flowers, might result in lower visitation to complex flowers (McCall and Primack 1992, Lázaro et al. 2013, Zhao et al. 2016). These contrasting effects of restricting access to floral resources have been observed in multiple studies, where some work has reported floral visitors favouring complex flowers (Stout et al. 1998) and others have found that floral visitors prefer flowers where resources can be accessed more easily (McCall and Primack 1992, Kunin and Iwasa 1996, Stout et al. 1998, Lázaro et al. 2013). The choices of floral visitors in communities where resources are available in a range of restrictions levels are likely contingent on the identity of competitor species (Stout et al. 1998), abundances of plant species (Kunin and Iwasa 1996), and degree of floral trait overlap between plant species (Hargreaves et al. 2009, Lázaro et al. 2013). Thus, plant species that restrict access to resources can potentially be at a competitive disadvantage under certain conditions, and this is likely contingent on the abundance of unrestricted resources offered by the co-flowering community, as well as the degree of floral trait overlap.

One way in which plants that offer pollen as primary reward can restrict access to pollen grains is through poricidal anthers (Buchmann 1983, van der Kooi et al. 2021). Some species of bees have evolved the capacity to produce vibrations (also known as floral vibrations or sonication) that facilitate the removal of pollen grains from poricidal anthers (Buchmann 1983, De Luca and Vallejo-Marín 2013). The interaction between plants with specialised floral morphologies, such as poricidal anthers, and the bee behaviour of deploying floral vibrations has given rise to the phenomenon of buzz pollination (Buchmann 1983, Vallejo-Marín 2019). During buzz-pollination, bees typically grasp the anthers with their mandibles, curl their bodies around the anthers, and then generate vibrations that result in pollen being released from the anthers through apical slits or pores (De Luca and Vallejo-Marín 2013). Using vibrations for pollen extraction is likely energetically expensive, as the production of floral vibrations by bees involves rapid contraction of the same thoracic muscles that power energetically costly wingbeat during flight (King and Buchmann 2003). During flight, these muscles consume as much as 100 times the energy than the resting metabolic rate (Dudley 2002). Floral vibrations have higher frequency and amplitude (velocity, acceleration, and displacement) than flight vibrations (Pritchard and Vallejo-Marín 2020) and, therefore, it is likely that floral vibrations are equally or more energetically costly as those produced during flight. Because of the energetic costs associated with vibratile pollen extraction, we might expect bees to favour more easily accessible pollen resources under certain circumstances. Buzz-pollination is prevalent among both plants (6-8% of all plant species across 65 families (Buchmann 1983) and bees (half of all bee species (Cardinal et al. 2018)), however, our understanding of how visitation to buzz-pollinated plants is influenced by the availability of unrestricted pollen resources in the surrounding plant community is limited.

Recent work has shown that competition for pollination services between buzz-pollinated individuals is prevalent (Mesquita-Neto et al. 2018, Soares et al. 2020). It is likely that these plant species also compete for pollinator services with non-poricidal taxa with unrestricted pollen resources. We hypothesise that due to the energetic and, potentially, learning costs associated with using vibrations for pollen extraction (Laverty 1980, Russell et al. 2016), visitation to plants with poricidal anthers should be reduced when flowers that do not restrict access to pollen are available in high abundances in a community. An alternative hypothesis is that because only a subset of floral visitors in a community can use vibrations to extract pollen, plants with poricidal anthers could potentially act as a private and reliable pollen resource to particular bee species, which could result in consistent visitation from these bees regardless of community context.

The Cape Floristic Region (CFR) of South Africa is well-suited for studying the effects of variation in co-flowering species composition on a focal plant species because it has sharp spatial and temporal changes in the composition of flowering communities (Cowling 1992, Simmons and Cowling 1996). Our study focuses on buzz-pollinated *Cyanella hyacinthoides* (Tecophilaeaceae) in the CFR. This species is widespread and thus co-occurs with a variety of other plant species, making it ideal to study the effects of co-flowering species composition on visitation. *Cyanella hyacinthoides* is a cormous geophyte endemic to the CFR that flowers during Austral spring (August to November) (Manning and Goldblatt 2012). It has light blue flowers with six poricidal anthers that vary in morphology (five smaller upper anthers and one larger lower anther) (Dulberger and Ornduff 1980). Plants from this species can present multiple inflorescences and each inflorescence can produce up to 15 flowers. Individual flowers can remain open for six or seven days if not pollinated (Dulberger and Ornduff 1980), but flowers close within a few hours after pollination has occurred (JEK, personal observation). Self-compatibility varies between populations, with two-thirds of assessed populations exhibiting complete self-incompatibility (Dulberger and Ornduff 1980), indicating that the persistence of this species mostly relies on successful pollinator-mediated reproduction.

Here, we contrast how visitation by bees that can successfully manipulate the buzz-pollinated *C. hyacinthoides* is influenced by the availability of more easily accessible resources, i.e., co-occurring plant species with simple floral morphologies. We predict that if *C. hyacinthoides* competes with flowers with unrestricted pollen for pollination services, there should be a reduction in the number of pollinator species and their visitation rates to *C. hyacinthoides* when the abundances of flowers with unrestricted pollen resources are high. Further, because floral trait overlap between co-flowering taxa can increase competition, we evaluated whether the pollinators of *C. hyacinthoides* were preferentially visiting flowers with particular floral traits. We predict that if bees favour particular floral traits, plant species with these traits might be more prominent competitors of *C. hyacinthoides.* Our study addresses the following questions: 1) How does the number of species visiting *C. hyacinthoides* varies when the availability of unrestricted rewards changes? 2) How do visitation rates to *C. hyacinthoides* vary when the availability of unrestricted pollen resources changes? 3) Which floral traits modulate visitation by the pollinators *of C. hyacinthoides*?

## Methods

### Sampling sites

We conducted our study in the Pakhuis Pass region of the northern Cederberg mountain range, situated in the west of the CFR in September-October 2019 (Fig. 1). This winter-rainfall area receives an annual rainfall of 270 ± 60 mm (mean ± SD; Pauw and Stanway (2015)). Most of the flowering in this region occurs during late-winter/early-spring (late-July to early-September), and flowering quickly ends when temperatures increase (Pauw and Stanway 2015). Buzz-pollinated species in this region generally start flowering in September, that is, after most other flowering has ended, and flowering continues throughout summer (Manning and Goldblatt 2012). The Cederberg region is home to multiple buzz-pollinated species, including *Cyanella hyacinthoides* (Tecophilaeaceae), *C. orchidiformis, C. alba, Ixia scillaris* (Iridaceae), *Solanum tomentosum* (Solanaceae), *Chironia linoides* (Gentianaceae), *C. baccifera,* and *Roridula dentata* (Roridulaceae), amongst others. For this study, we focused on communities containing *Cyanella hyacinthoides* as it is widespread and occurs among a wide variety of co-flowering species.

**Figure 1.**
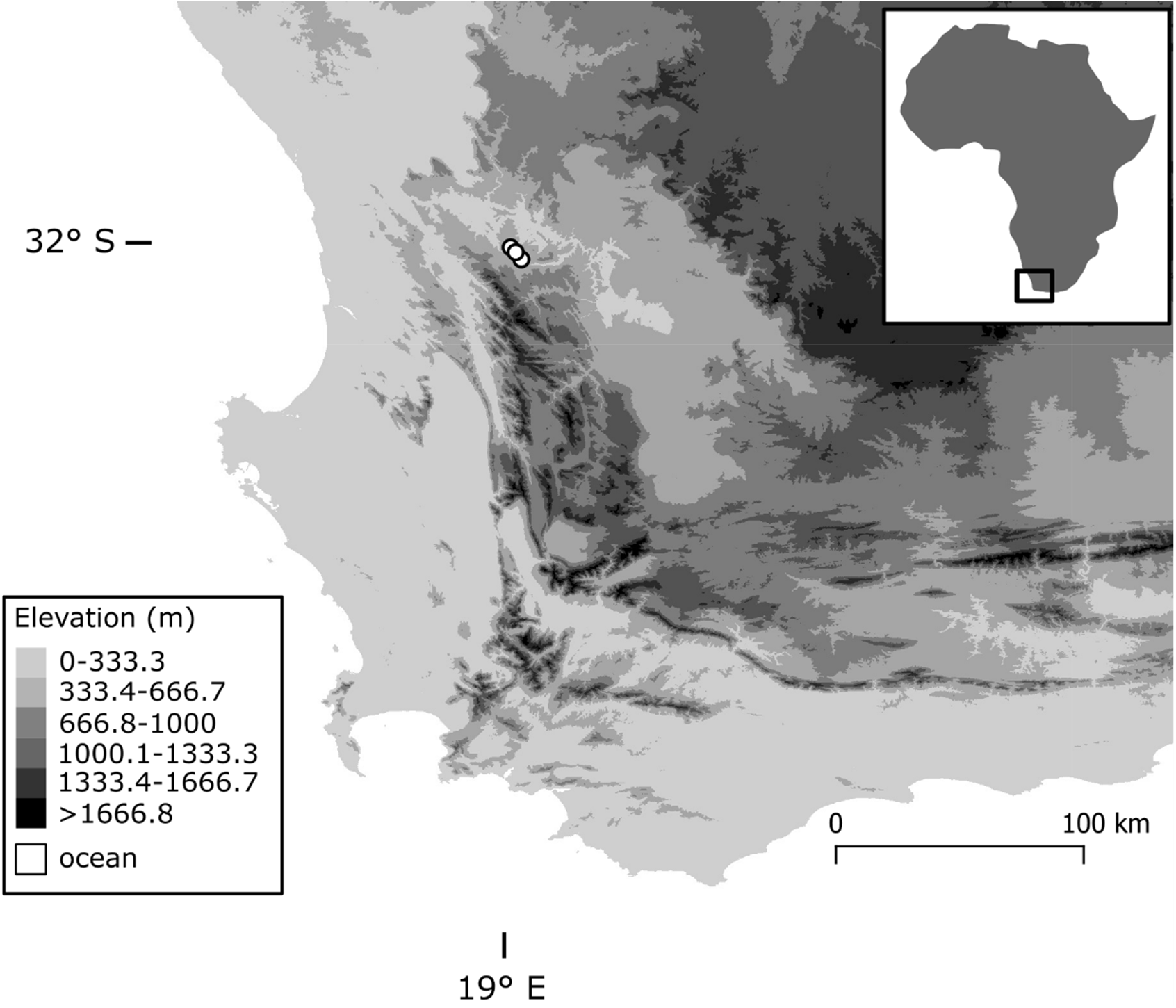
Elevation map of south-western South Africa, which primarily contains the hyperdiverse Cape Floristic Region. The sampling sites were located in the northern Cederberg mountain range, indicated by white circles.

We identified three sites where *C. hyacinthoides* was abundant. The sites present a shrubland vegetation that is classified as Rainshadow Valley Karoo (Mucina et al. 2006), located in the transition between fynbos and the drier inland vegetation (Fig. 2). Each site covered approximately 100 m x 100 m, and the sites were separated by 4-10 km. We exploited the CFR’s tendency to sharp temporal turnover in community composition, and we specifically chose sites where (A) few individuals of *C. hyacinthoides* recently started flowering and many individuals had not yet started flowering (i.e., *C. hyacinthoides* would continue to flower for at least two to three more weeks), (B) multiple plant species with unrestricted pollen resources were at a late stage of flowering (i.e., few buds and many seed pods present, and flowering would end within a week), and (C) where multiple plant species with unrestricted pollen resources were still in bud stage (i.e., would start flowering in about a week). These communities thus showed a change in *C. hyacinthoides* flower abundances over a short timeframe as more individuals start flowering, as well as a change in the composition of other plant species as the late-stage flowering species (B) end their flowering and the bud-stage flowers (C) initiate flowering (see *Results* for details on plant species turnover). By sampling each of these three sites twice, we were able to observe the pollination interactions of *C. hyacinthoides* in different community contexts, whilst controlling for environmental and site-specific factors. To verify that the communities showed sufficient spatial and temporal variation in flower composition, we quantified plant turnover by calculating beta diversity (Horn similarity, following Jost 2007).

**Figure 2.**
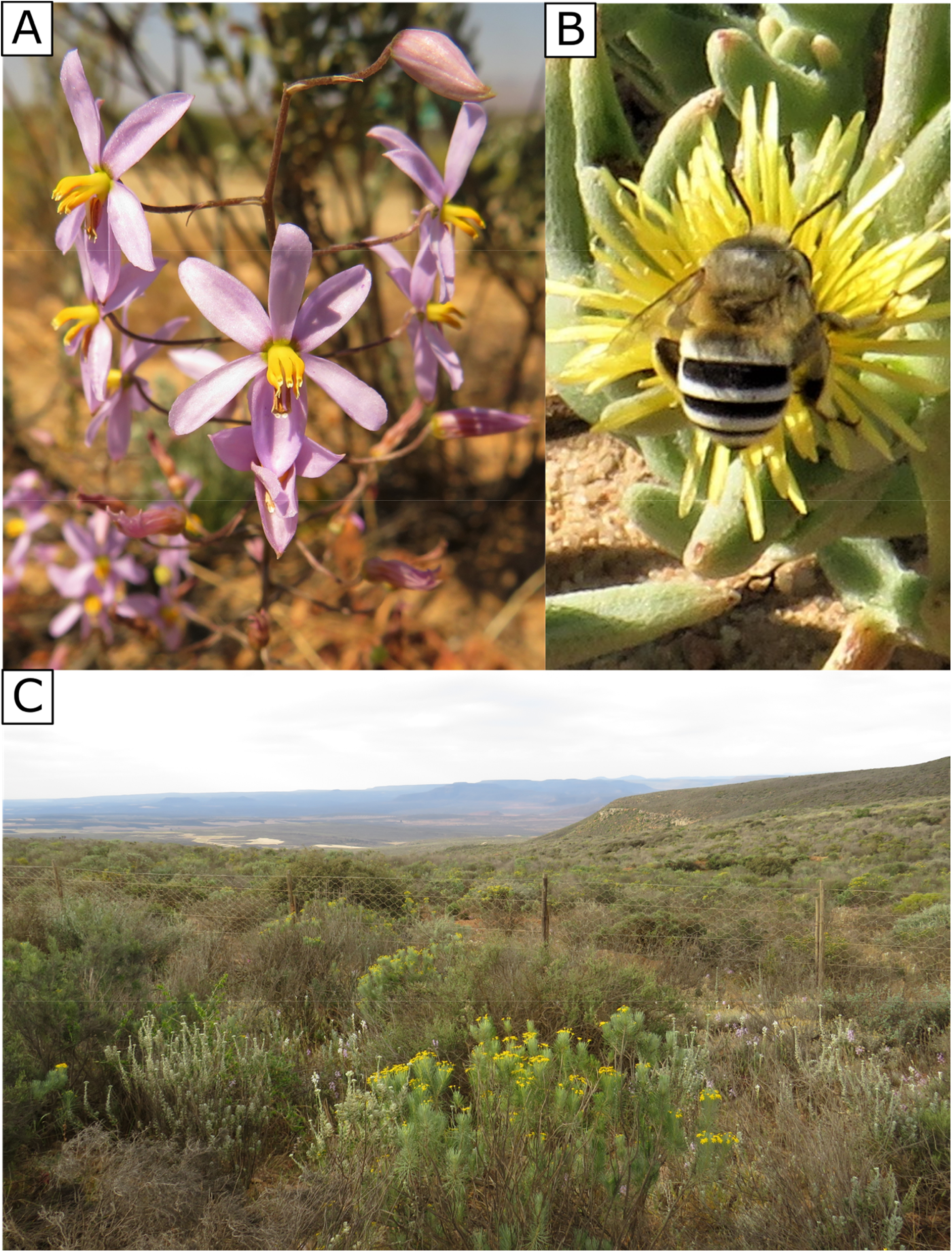
*Cyanella hyacinthoides*, its main pollinator, and the vegetation type of the sites are shown. (A) *C. hyacinthoides* is a cormous perennial that presents flowers with poricidal anthers on multiple inflorescences. Flowers are approximately 16 mm in diameter across the longest axis. (B) *Amegilla* cf. *niveata* was the most frequent visitor to *C. hyacinthoides* in these communities. In this photo, a female is drinking nectar from a *Cleretum* flower (flower approximately 16 mm in diameter). (C) The sites were situated in Rainshadow Valley Karoo shrubland. The shrub in the foreground with yellow flowers is approximately 80 cm tall. Photos: JEK.

**Figure 3.**
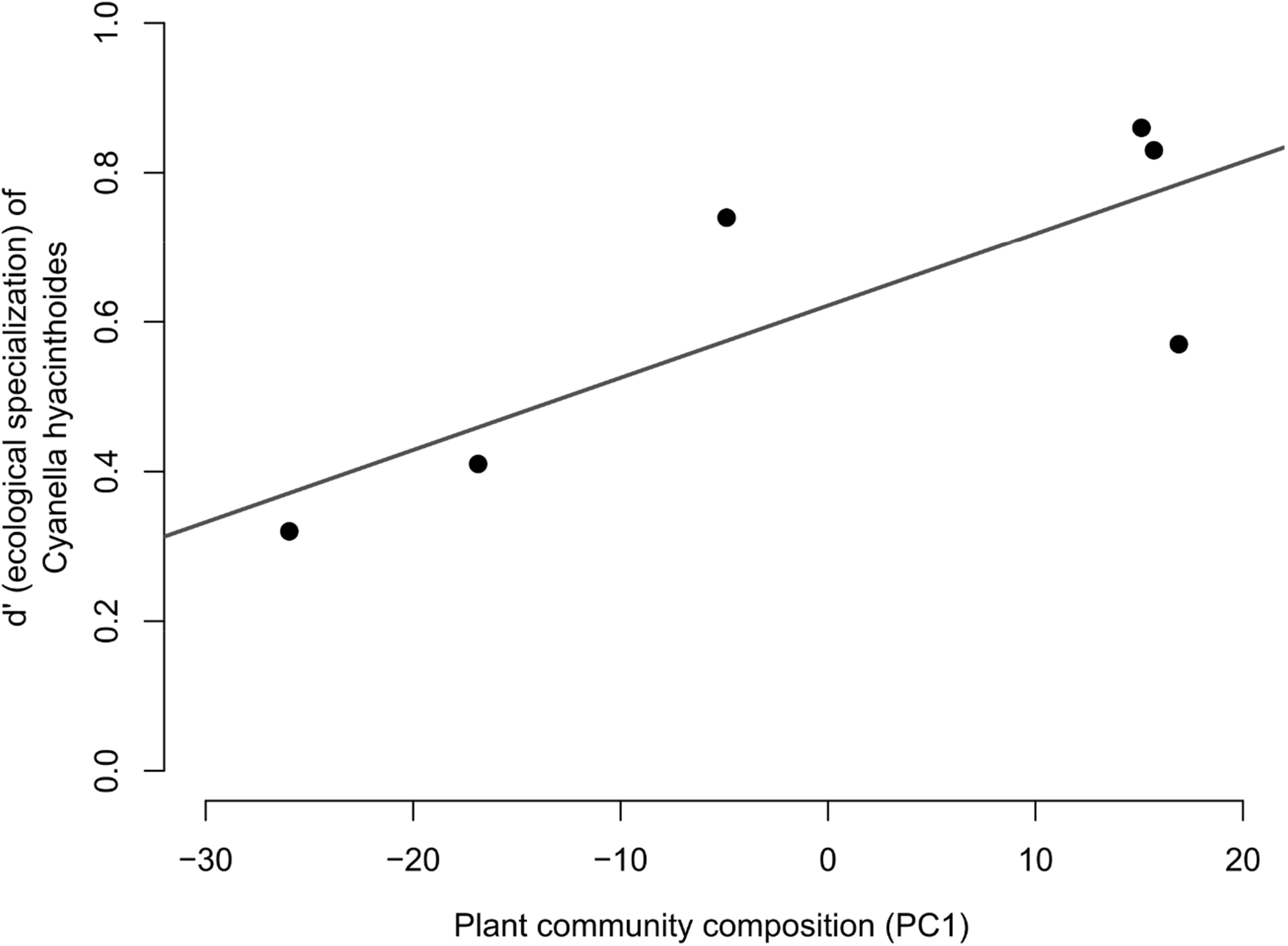
Ecological specialization of *Cyanella hyacinthiodes* in its pollination interactions (measured as d’) was associated with the composition of pollen-offering plant species. This specialization metric indicates whether a pollinator species is selectively visiting a plant species relative to other available floral resources (1 = high selectivity, 0 = low selectivity). Plant community composition is represented by the first principle component of a PCA conducted on pollen-offering plant species composition, and high values are associated with high *C. hyacinthoides* abundances and low values are associated with high *Euryops tennuissisimus* (Asteraceae) abundances.

### Pollinator observations

For each of the three sites, two observers recorded interactions over three to five days, depending on the weather. Each site was sampled a second time, approximately nine days after the first sampling session ended. This resulted in the six data sets that we consider as six communities (see *Results* for turnover in plant composition between sampling sessions). The first site was sampled from 17-22 September 2019, and resampled 1-4 October 2019. The second site was sampled 22-25 September 2019, and resampled 4-6 October 2019. The third site was sampled 25-28 September 2019, and resampled 8-12 October 2019.

Floral visitation observations commenced when bee activity started, that is, after 06h00, depending on the weather. If the temperature exceeded 30°C and bee activity decreased, observations were halted and resumed later in the day. Interactions between bees and flowers were recorded in 20-minute intervals, and multiple plant species were observed simultaneously. The number of flowers per species that were observed per interval was noted. In total, interactions were observed for 607 intervals, resulting in 202.3 observation hours. Interaction strengths were calculated as the number of visits per flower per 20-minute interval multiplied by 1000 and rounded to create integers. We did this as some of our analyses required integers as input.

Our analyses (described below) relied on the interaction frequencies of vibrating bees to all plant species in the community from which they were observed to collect pollen. We thus identified the vibrating bees to genus or species level using the keys by Eardley (1994) and Eardley & Brooks (1989) The plant species that were visited by the vibrating bees were identified to species level using Manning and Goldblatt (2012). Further, some of our analyses required the total number of visits made by all bee species to all pollen-offering plant species in each community, particularly for the calculation of d’ and link temperature (see below). For these calculations, only the sum of all the interactions in the community is relevant and the identities of bee and plant species do not matter. We thus did not identify these additional bee and plant species to species level, however, we identified these additional bee species as morphospecies in the field (through capture, behaviour, and photographs), and we identified plant species to genus level, to assist with our observations in the field.

### 1) *How does the number of species visiting* C. hyacinthoides *change when the availability of unrestricted resources changes?*

The number of bee species visiting a plant species (i.e., ecological pollination specialization – *sensu* Armbruster (2017)) can be described by multiple metrics, and we calculated three metrics that captured this within each community (Table 1). The first metric is *interaction partner richness*, which measures the raw number of bee species that visits *C. hyacinthoides*. For instance, if five species visit *C. hyacinthoides*, then the interaction partner richness would be 5. The second metric is *interaction partner diversity*, calculated using Hill numbers of the Shannon diversity index (Jost 2007). This metric calculates the number of bee species that visits *C. hyacinthoides* and weights it with the interaction frequency of each pollinator species, thus accounting for interaction evenness (Kemp et al. 2019). Thus, if *C. hyacinthoides* is visited by many insect species, but only a few of these species have high visitation rates to *C. hyacinthoides*, this metric will indicate that *C. hyacinthoides* is effectively visited by few pollinator species. For instance, if one pollinator species makes ten visits to *C. hyacinthoides* and four species each make one visit, the interaction partner diversity would be 2.70. This metric thus adjusts for uneven visitation by pollinators. Finally, we calculated the *specialization index d’* which indicates the specialization of *C. hyacinthoides* in relation to the availability of bee species in the community (Blüthgen et al. 2006). This metric ranges from 0 to 1, where high values indicate that a plant species is selectively visited by few insect species, and low values indicate that a plant species is either visited by many insect species or by the common insect species in the community. Ultimately, this metric shows how strongly a plant species deviates from a random sampling of the available pollinators. For instance, if a bee species that visits *C. hyacinthoides* also visits all plant species in the community at similar frequencies, then *C. hyacinthoides* will have a low d’ value because it is not selectively visited by the bee species. However, if a bee species makes many visits to *C. hyacinthoides* and few visits to other plant species in the community, then *C. hyacinthoides* will have a high d’ value because the bee species is selectively visiting it rather than other available plant species.

**Table 1.**
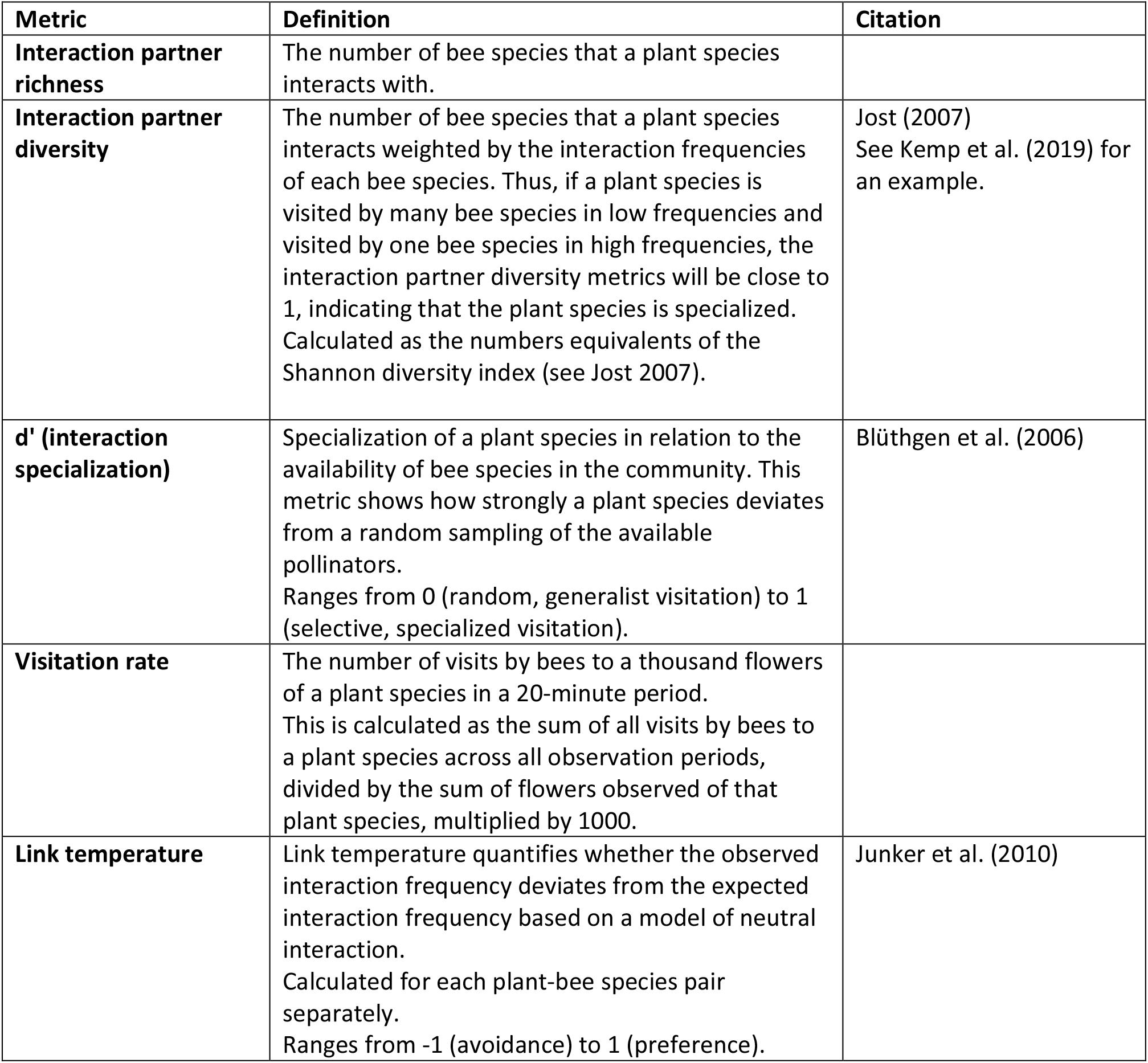
Summary of the metrics used to assess the variation the pollination interactions of *Cyanella hyacinthoides*.

| Metric | Definition | Citation |
| --- | --- | --- |
| **Interaction partner richness** | The number of bee species that a plant species interacts with. |  |
| **Interaction partner diversity** | The number of bee species that a plant species interacts weighted by the interaction frequencies of each bee species. Thus, if a plant species is visited by many bee species in low frequencies and visited by one bee species in high frequencies, the interaction partner diversity metrics will be close to 1, indicating that the plant species is specialized. Calculated as the numbers equivalents of the Shannon diversity index (see Jost 2007). | Jost (2007)<br>See Kemp et al. (2019) for an example. |
| **d' (interaction specialization)** | Specialization of a plant species in relation to the availability of bee species in the community. This metric shows how strongly a plant species deviates from a random sampling of the available pollinators.<br>Ranges from 0 (random, generalist visitation) to 1 (selective, specialized visitation). | Blüthgen et al. (2006) |
| **Visitation rate** | The number of visits by bees to a thousand flowers of a plant species in a 20-minute period.<br>This is calculated as the sum of all visits by bees to a plant species across all observation periods, divided by the sum of flowers observed of that plant species, multiplied by 1000. |  |
| **Link temperature** | Link temperature quantifies whether the observed interaction frequency deviates from the expected interaction frequency based on a model of neutral interaction.<br>Calculated for each plant-bee species pair separately.<br>Ranges from -1 (avoidance) to 1 (preference). | Junker et al. (2010) |

To assess whether ecological specialization of *C. hyacinthoides*, as measured by the three metrics created above, changes when the surrounding community composition changes, we first reduced community composition into a single variable by conducting a principal component analysis (PCA) and keeping the first principal component. Because we are interested in plant species that potentially act as competitors to *C. hyacinthoides*, we only included plant species in the PCA that (A) received 10 or more visits per 1000 flowers in any community from bees that use vibrations to extract pollen from *C. hyacinthoides* (hereafter called vibrating bees) and (B) was observed to have pollen collected by vibrating bees in any community. Five plant species met this requirement and were included in the PCA: *Convolvulus capensis* (Convulvulaceae), *Cyanella hyacinthoides* (Tecophilaeaceae), *Roepera morgsana* (Zygophyllaceae), *Ornithogalum thyrsoides* (Hyacinthaceae), and *Euryops tenuissimus* (Asteraceae). Although the vibrating bees occasionally collected nectar from some of these plant species (*C. convolvulus and R. morgsana*), we still consider them as potential competitors for pollen collection by vibrating bees. We conducted a PCA of the plant densities per m^2^ for each community using the ‘prcomp’ function in R (R core team, 2020). The first principal component explained 95% of variation, and high positive values along this axis was associated with high *C. hyacinthoides* (loading PC1: 0.15) abundances and negative values were associated with high *E. tenuissimus* (loading PC1: −0.98) abundances. The other three species contributed very little to the first principal component (loading PC1: *O. thyrsoides*: 0.07; *R. morgsana*: 0.04; *C. capensis*: −0.02). We then used this first principle component as proxy for community composition in our model. Next, we calculated the total number of visits that vibrating bees made to all pollen-offering plants in a community. This was calculated by multiplying the visits per flower with the flower density per m^2^, and then multiplying this with 10,000 m^2^ (approximate community size), giving the number of visits each vibrating bee species made to each plant species. Visits were summed across plant species to give the total number of visits made by these bee species within a community, and we used this as proxy for the abundances of vibrating bees.

We conducted three models with the specialization metrics (i.e., interaction partner richness, interaction partner diversity, and d’) as respective response variables. Vibrating bee abundances and community composition (as represented by PC1 from the PCA above) as calculated above were used as explanatory variables. Specifically, Poisson regression was used to test the influence of vibrating bee abundances (natural log-transformed to improve model fit) and community composition on interaction partner richness. Further, we conducted two linear regressions to assess the influence of vibrating bee abundances (natural log-transformed to improve model fit) and community composition on interaction partner diversity and *d’* respectively.

### 2) *How do visitation rates to* C. hyacinthoides *change when the availability of unrestricted resources changes?*

To assess whether the surrounding plant community influences visitation rates to *C. hyacinthoides,* we conducted a negative binomial regression, to accommodate overdispersion of the data, with the number of visits per thousand *C. hyacinthoides* flowers by vibrating bees as response variable. We used the first principal component of plant composition (from the previous section) and the bee abundances calculated above (natural log-transformed to improve model fit) as predictor variables.

### 3) *Which floral traits modulate visitation by the pollinators of* C. hyacinthoides*?*

For each plant species, we measured the plant height and flower diameter (along its longest dimension for asymmetric flowers). For this, we measured a median of 30 flowers per species. Additionally, we recorded flower colour by measuring reflectance spectra indoors at a 45° angle to the petal surface with an OceanOptics USB4000 Spectrometer (Ocean Optics, Dunedin, FL, USA) calibrated with a diffuse reflectance WS-2 white standard. Multiple measurements were taken per species (mean = 14, range = 6-30), and these were averaged to obtain a single spectrogram per species. We modelled these spectra in honeybee vision using Chittka’s hexagon model (Chittka 1992), assuming a D65 illumination and a standard green background, in the ‘pavo’ package (Maia et al. 2016) in R. We chose the honeybee visual model because the specifics of the photoreceptors of most of the bee species in this study are unknown, except for honeybees. Plant species were subsequently grouped into six categories in the hexagon (i.e., blue, UV-blue, UV, UV-green, green, and blue-green) based on the relative excitations of the three types of bee photoreceptor (UV, blue and green). However, none of the plant species had bee-blue petals, and thus all plant species were assigned to one of the other five groups.

To assess whether floral traits are associated with visitation by vibrating bees, we calculated link temperature (Junker et al. 2010) for all of the plant species in each network. Link temperature quantifies whether the observed interaction frequency deviates from the expected interaction frequency based on a model of neutral interaction. Link temperature is calculated for each insect species separately, and we thus calculated the link temperature between each of the three vibrating bee species and all of the plant species in our communities. Link temperature (*T*_*ij*_) is calculated as:

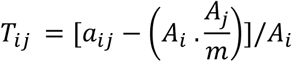

Where *a*_*ij*_ is the observed number of interaction between bee species *i* and plant species *j*. *A*_*i*_ represents the total number of visits made by bee species *i* to all plant species in the community, and *A*_*j*_ represents the total number of visits received by plant species *j* from all bee species. The grand total of all interactions recorded between all plant species and all bee species in a community is given by *m*. Link temperature ranges from −1 to 1, where −1 indicates that a pollinator is avoiding a plant species and 1 indicates a pollinator is favouring a plant species (see Junker et al. 2010).

To test whether bees favour or avoid plant species based on visual traits, we implemented a linear mixed effect model using link temperature as response variable using the ‘lme4’ package in R (Bates et al. 2015). We conducted stepwise backward model selection to determine which predictors to include in the final model (‘step’ function in ‘lmerTest’ (Kuznetsova et al. 2017). The model selection procedure included plant height, flower diameter, flower colour group in bee space, and vibrating bee species as predictor variables. Because some plant species were present in multiple communities, we included plant species as random factor. Model selection using Satterthwaite’s type III ANOVA indicated that only colour group should be included in the final mixed effect model, with plant species as random effect. To test for differences between the effects of the different colour groups on link temperature in the final model, we performed t-tests using Satterthwaite’s method.

## Results

The six communities studied consisted of 26 plant species, of which 20 species offered pollen resources to bees (determined from observing pollinators on flowers). Communities showed both spatial and temporal turnover in plant species, based on flower densities recorded in 25-30 random 4 m^2^ plots at each site. In the first sampling session, the three communities showed 50.57% (sd = 13.76) similarity in plant species (Horn similarity - Jost, 2007), and in the second session, communities showed 62.00% (sd = 20.01) similarity in plant species composition. Between sampling sessions (i.e., temporal similarity), communities showed 54.48 % (sd = 16.71) similarity in plant species composition. The six communities thus showed sufficient turnover in plant composition for our purposes.

We recorded visits from 180 insect morphospecies, of which 66 morphospecies were bees. Only three bee species visited *C. hyacinthoides* all of which used vibrations to extract pollen from this species. Two of these bee species were present in all six communities, and only one of these visited *C. hyacinthoides* in all six communities.

The number of bee species visiting *C. hyacinthoides*, measured as interaction partner richness and diversity, was not influenced by the availability of other pollen (Table 2). However, the specialization index d’ was positively associated with the first principle component of plant community composition (Table 2, Fig. 5), indicating that bee species selectively visited *C. hyacinthoides* in communities with high *C. hyacinthoides* abundances.

**Table 2.** Association between various measures of ecological specialization of *C. hyacinthoides* and community composition (plant and insect). Plant community composition is represented by the first principle component of a PCA conducted on the abundances of five plant species that are visited by *C. hyacinthoides*’s pollinators. Vibrating bee abundances are the abundances of *C. hyacinthoides*’s pollinators. The association between the response and predictor variables were tested using a Poisson regression (for interaction partner richness) and linear regressions (for interaction partner diversity and d’). Significant associations at p < 0.05 are indicated in bold.

| Response | Predictor | estimate | z or t | p |
| --- | --- | --- | --- | --- |
| Interaction partner richness | Plant composition | 0.0127 | 0.622 | 0.53 |
|  | Vibrating bee abundance (log-transformed) | -0.0125 | -0.098 | 0.92 |
| Interaction partner diversity | Plant composition | 0.0140 | 0.963 | 0.41 |
|  | Vibrating bee abundance (log-transformed) | -0.3092 | -0.920 | 0.43 |
| d' | **Plant composition** | **0.0138** | **4.557** | **0.02** |
|  | Vibrating bee abundance (log-transformed) | 0.1651 | 2.355 | 0.10 |

The number of visits to *C. hyacinthoides* flowers were positively associated with the first principle component of plant community composition (GLM: estimate = 0.030, z = 2.336, p = 0.02, Fig. 4A), which shows that visitation rate to *C. hyacinthoides* increased with an increase in *C. hyacinthoides* abundances. Further, visitation to *C. hyacinthoides* was positively associated with the abundances of vibrating bees (GLM: estimate = 1.438, z = 4.773, p < 0.001, Fig. 4A).

**Figure 4.**
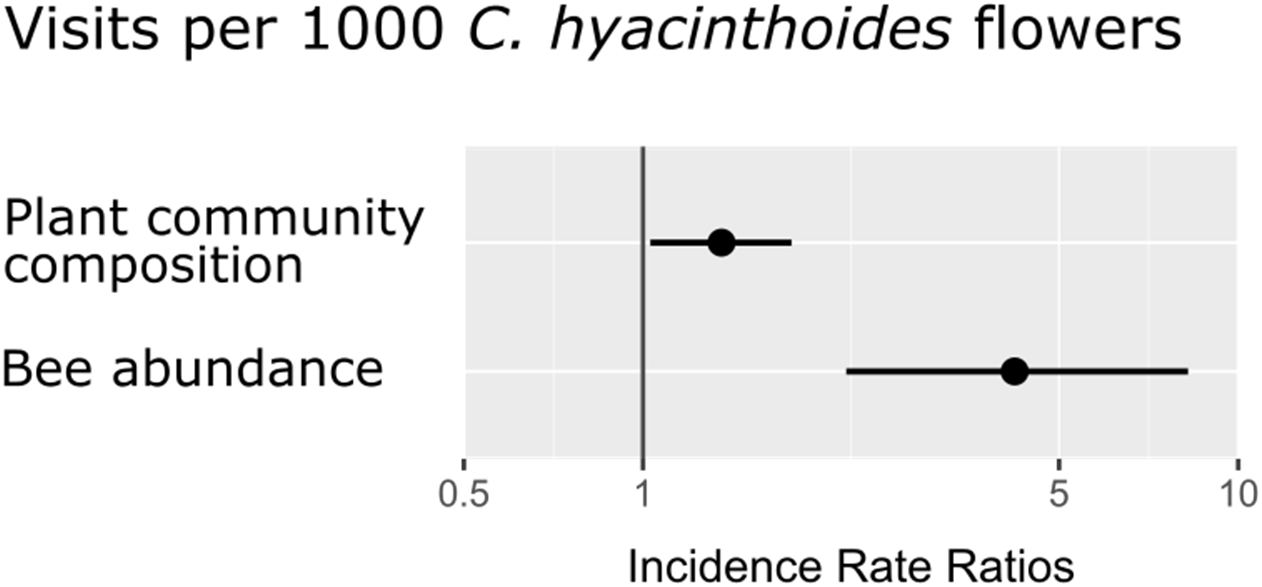
Results from negative binomial general linear model on the influence of plant community composition and bee abundances on the visitation rates (i.e., visits per 1000 flowers) to *Cyanella hyacinthoides* across six communities. Plant community composition is presented by the first principle component from a PCA conducted on plant abundances, and the total number of visits made to all pollen-offering plants within a community by vibrating bees were used as a proxy for vibrating bee abundances. Incidence rate ratios larger than 1 indicate a positive association between the response and predictor variables, and values smaller than 1 indicate a negative assocaition. Both predictor variables were significantly positively associated with the visitation rates to *C. hyacinthoides.*

Our stepwise model selection procedure showed that colour group was the only important predictor of link temperature (Table 3), and plant height, pollinator species, and flower diameter were thus excluded from the final model. Linear mixed effect modelling showed that link temperature was lower for flowers that fall into the green and UV-blue hexagon sections compared to flowers classed as blue-green (Fig. 5, Table 3), suggesting that vibrating bees avoided green and UV-blue flowers compared to blue-green flowers. The median link temperature for bee-blue-green flowers was 0.02 (sd = 0.35), suggesting that these bees preferentially visited flowers in this colour category (positive values indicate preference and negative values indicate avoidance – Table 1).

**Table 3.**
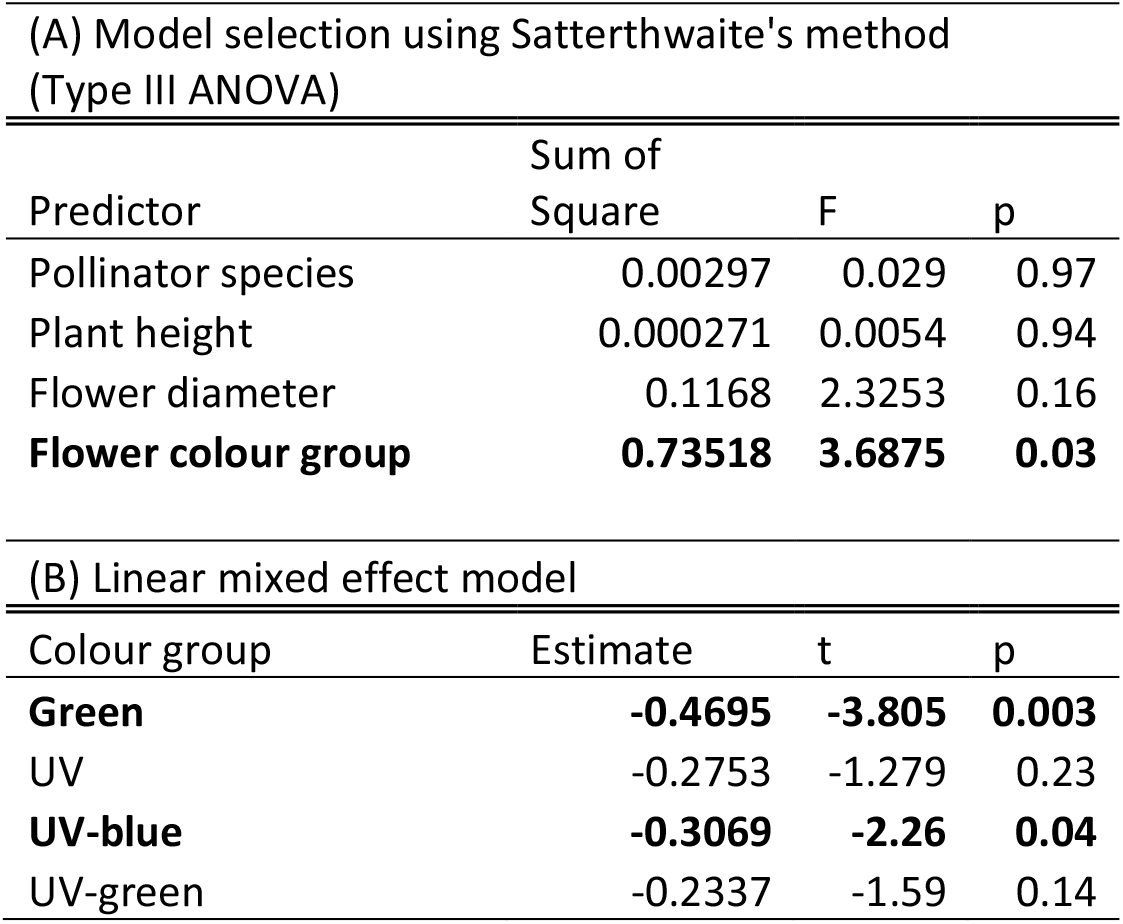
Backward model selection and subsequent linear mixed effect model results for features predicting link temperature of bees that use vibrations for pollen extraction. (A) Backward model selection was performed using the ‘step’ function in the lmerTest package (Kuznetsova et al. 2017) with the link temperature of bees as response variable, the four predictor variables listed in the table below as predictor variables, and plant species as random factor. Model selection was based on Satterthwaite’s Type III ANOVA, and flower colour group (i.e., flower colour category in the bee hexagon visual model) was the only important predictor of link temperature of bees that use vibrations for pollen extraction. (B) We subsequently performed a linear mixed effect model with the link temperature of bees as response variable, flower colour category as predictor variable, and plant species as random factor. Pairwise differences between groups were determined through t-tests using Satterthwaite’s method. The colour group blue-green is absorbed in the model. P-values lower than 0.05 are indicated in bold.

| Predictor | Sum of Square | F | p |
| --- | --- | --- | --- |
| Pollinator species | 0.00297 | 0.029 | 0.97 |
| Plant height | 0.000271 | 0.0054 | 0.94 |
| Flower diameter | 0.1168 | 2.3253 | 0.16 |
| **Flower colour group** | **0.73518** | **3.6875** | **0.03** |

(B) Linear mixed effect model
| Colour group | Estimate | t | p |
| --- | --- | --- | --- |
| **Green** | **-0.4695** | **-3.805** | **0.003** |
| UV | -0.2753 | -1.279 | 0.23 |
| **UV-blue** | **-0.3069** | **-2.26** | **0.04** |
| UV-green | -0.2337 | -1.59 | 0.14 |

**Figure 5.**
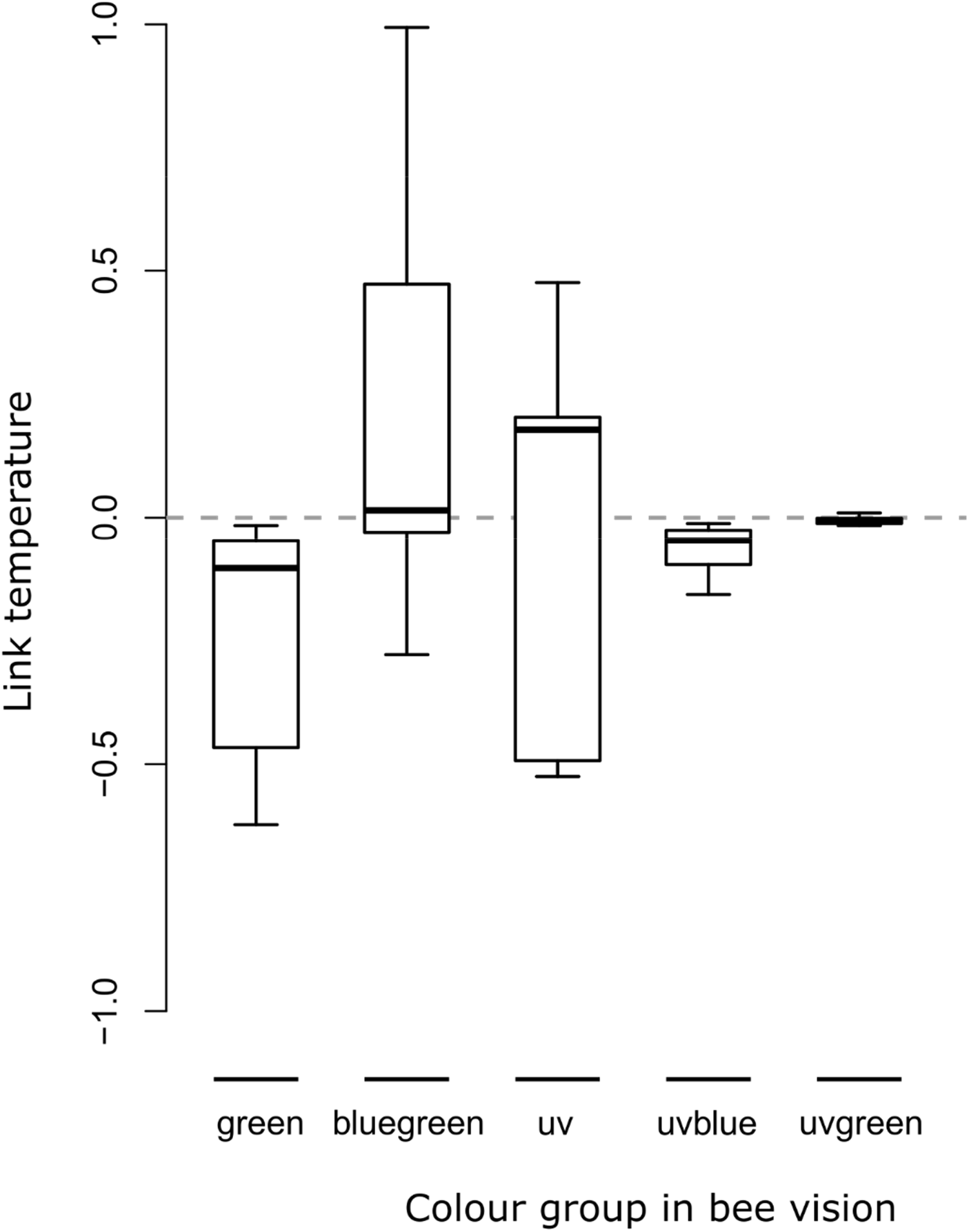
Differences in the link temperature between flowers of difference colour categories (as perceived by bees). High values of link temperature indicate that bees are preferentially visiting flowers of a particular colour group, and low values indicate bees are avoiding a particular colour group. Link temperature was calculated for all pollen-offering plant species and the pollinators of *Cyanella hyacinthoides*.

## Discussion

We investigated the variation in pollination interactions (i.e., number of pollinator species and their visitation rates) of the buzz-pollinated *Cyanella hyacinthoides* when it occurred in different co-flowering communities. Although the number of bee species that visited *C. hyacinthoides* was not influenced by the availability of more easily accessible pollen resources, the visitation rates of these bees were influenced by the co-flowering community composition. Specifically, *Cyanella hyacinthoides* received more visits per flower when many conspecifics were present in a community, and fewer visits per flower when many other pollen resources were available. Further, we show that bees exhibited non-random visitation to flowers within these communities and avoided or preferred flowers with certain petal colours. Our results support the hypothesis that buzz-pollinated flowers might be at a competitive disadvantage when more easily accessible pollen resources are abundant, particularly when the competitor species have similar floral traits.

### Cyanella hyacinthoides *is visited by few bee species*

*Cyanella hyacinthoides* flowers were visited and vibrated by a small subset of the available bee species (4.5% of morphospecies). Only three of the 66 bee morphospecies were observed to use vibrations to extract pollen from *C. hyacinthoides*, with most of these visits made by *Amegilla* cf. *niveata*. Our results showing pollination specialization in *C. hyacinthoides* aligns with findings from other buzz-pollinated systems, where only a subset of bees in a community visited buzz-pollinated taxa (Goldblatt et al. 2000, Mesquita-Neto et al. 2018, Soares et al. 2020). Interestingly, Mesquita-Neto et al. (2018) found that different buzz-pollinated plant species within a community in Brazil were visited by different subsets of vibrating bees, which is indicative of pollination niche partitioning among buzz-pollinated plant species within a community. Each plant species was visited by 1 to 15 bee species that successfully vibrated flowers (of the 55 available bee species). The niche partitioning amongst buzz-pollinated species that Mesquita-Neto et al. (2018) observed suggests that floral traits other than poricidal anthers might be important in enabling or limiting visitation to buzz-pollinated flowers. Our results show that in our system, the colour of the petals mediates pollination interactions in co-flowering communities. The three bee species that visited and vibrated *C. hyacinthoides* visited other plant species in the community to collect pollen, but they visited only a small subset of the unrestricted pollen sources, suggesting that they are not generalist pollen foragers. Further study is required to determine which floral traits mediate pollination interactions in other buzz-pollinated taxa, and whether buzz-pollinated plants are frequently visited by pollen-specialist bees.

### High availability of other pollen sources is associated with less buzz pollination

Our results demonstrate a density dependent effect of visitation rates by vibrating bees to *C. hyacinthoides.* In communities where flowers with easily accessible pollen were abundant, *C. hyacinthoides* received fewer visits per flower than in communities where *C. hyacinthoides* occurred in high abundances. Further, the specialization index d’ was related to community composition, which indicates that in communities with high *C. hyacinthoides* abundances, pollinators were preferentially visiting *C. hyacinthoides* rather than other available plant species. Our results are supported by previous studies on flowers that are not poricidal but require complex handling behaviours. For instance, Stout et al (1998) found that some species of complex flowers were at a competitive disadvantage in the presence of simple flowers with easily accessible resources. However, this was dependent on the identity of both the complex and simple flowered species, which suggests that additional floral traits are important in the foraging decisions of bees. In contrast to our results, Lázaro et al. (2013) showed that visitation to complex flowers did not change with heterospecific flower density, but was rather related to pollinator abundances. Further, Lázaro et al (2013) showed that seed set increased with an increase in both con- and heterospecific flower densities, suggesting facilitative interactions were prevalent among plant species, in contrast to the competitive interactions we observed in our study system. One potential reason for the contrasting results might be due to the identity of co-flowering plant species and the rewards they offer (Stout et al. 1998). Although buzz-pollinated species likely compete with other pollen-offering flowers, the presence of nectar-offering species in a community will potentially have facilitative effects on visitation to buzz-pollinated flowers. The availability of nectar sources in a community might be particularly important for bees that use vibratile pollen extraction which requires high energy consumption (Pritchard and Vallejo-Marín 2020), and we might potentially expect the co-occurrence of nectar sources to be a prerequisite for the occurrence of buzz-pollinated species within a community, although this has not yet been investigated.

The reduced visitation rates to buzz-pollinated flowers when unrestricted pollen resources are abundant can likely be attributed to the metabolic and/or learning costs associated with vibratile pollen extraction. The metabolic costs associated with vibratile pollen extraction is potentially more than 100 times as costly as resting metabolic rates (Dudley 2002, Pritchard and Vallejo-Marín 2020), and this might deter bees from visitation to buzz-pollinated flowers when unrestricted pollen resources are readily available in a community. However, buzz-pollinated flowers have been suggested to produce higher quality pollen than non-buzz-pollinated plants (Roulston et al. 2000), and this might incentivise bees to learn to effectively manipulate them when they occur in sufficient abundances. Further, the time required to learn to extract pollen from complex flowers might deter pollinators from visiting buzz pollinated flowers. Bees generally take longer to learn to extract resources from flowers that require complex handling behaviours than those that do not (Gegear and Laverty 1998), but bees can remember flower-handling techniques for long time periods (Keasar et al. 1996, Chittka and Thomson 1997). Thus, if simple and complex flowers contain similar resources and are equally abundant, it is initially more costly for bees to visit flowers that require complex handling behaviours until these behaviours can be performed efficiently (Krishna and Keasar 2018). The low visitation rates to *C. hyacinthoides* when their relative abundances are low might thus potentially be attributed to costs associated with learning to handle a rare and complex flower type.

Our results have important implications for the reproductive ecology of *C. hyacinthoides* and other buzz-pollinated plant species, and buzz-pollinated plant species could potentially suffer reduced fitness in communities where they occur in low abundances. In non-poricidal plant species, high abundances of co-flowering conspecifics have been shown to increase pollinator visitation (Rathcke 1983, Moeller 2004), as well as pollen removal and deposition (Duffy and Stout 2011), due to the increased size of the floral display. Although an increase in plant density can increase per-flower visitation rates (e.g., Duffy and Stout, 2011), this will likely saturate at high plant densities due to pollinator limitation, and visitation rates are likely to decrease at higher plant densities. Our results contrast Stout et al. (1998) and Johnson et al. (2012) who showed that complex flowers received more visits per flower when they occurred in low densities than high densities. However, our sampling design did not include communities with extremely high flower densities (densities ranged from 1.08 to 11.23 *C. hyacinthoides* flowers per m^2^, and 0.15 – 2.06 *C. hyacinthoides* plants per m^2^), and thus we cannot rule out that the visitation rates might decrease at higher densities (e.g., Zimmerman (1980). Our results also show that visitation rates to *C. hyacinthoides* flowers is higher when vibrating bees make more visits at the community-level (i.e., our proxy for vibrating bee abundance). This suggests that the timing of flowering in *C. hyacinthoides* might be associated with the emergence of particular bee taxa, and *C. hyacinthoides* will likely suffer reduced fitness when flowering when few vibrating bees are available.

### Bees preferred and avoided flowers with certain colours

Our understanding of the motivation of bees to visit poricidal flowers within a community of flowering plants remains limited. We show that these interactions are partly mediated by the visual signals of *C. hyacinthoides* in this system. The pollinators of *C. hyacinthoides* visited plant species with bee-blue-green reflective petals and avoided those that primarily reflect bee-UV-blue or bee-green. In contrast to our results, previous work has shown that some bee species have innate preferences for bee-UV-blue (Giurfa et al. 1995) and for bee-green (Giurfa et al. 1995). Further, although *C. hyacinthoides* flowers reflected bee-blue-green, eight of the 26 species reflected this colour, and thus *C. hyacinthoides* did not provide a unique colour signal. We observed, however, that *C. hyacinthoides* flowers emitted a strong scent, and scent might be an important mediator in these buzz-pollination interactions, similar to what has been shown for other buzz-pollinated taxa (Solís-Montero et al. 2018, Vega-Polanco et al. 2020).

### Conclusions and Future Directions

Our work represents one of the first studies on the community ecology of buzz-pollination interactions, and we show that the co-flowering community of pollen-offering species influenced visitation to a buzz-pollinated species. Bees preferentially used vibratile pollen extraction when flowers with poricidal anthers occurred in high abundances, suggesting that it might be costly for bees to seek them out among other species when such flowers are rare. A promising avenue for future work encompasses determining whether this cost relates to time spent flying between low-density flowers, the energy cost of using vibrations for pollen extraction, or whether cognitive constraints limit the number of flower handling behaviours bees perform on a foraging bout. We also show that flower colour is important in mediating pollination interactions in these communities, and future work should investigate whether pollen-offering plant species with the same flower colour have facilitative or competitive effects on buzz-pollination interactions. Although we only explored the influence of the co-flowering pollen-offering species, the availability of flowers that offer nectar resources will likely also influence buzz-pollination interactions. Bees collect both pollen and nectar resources, and if bees prefer collecting nectar from particular plant species, the presence of these species in a community or in a patch might facilitate visitation to flowers with poricidal anthers. Additionally, these nectar sources might be important in providing bees with the necessary sugar (i.e., energy) to sustain vibratile pollen extraction.

## Acknowledgements

The project was funded by the Royal Society of London and the Newton Fund through a Newton International Fellowship to JEK (NIF/R1/181685), and was partially supported by a research grant from the Leverhulme Trust (RPG-2018-235) to MVM.

